## Supplementary material for "An estimate of the deepest branches of the tree of life from ancient vertically-evolving genes": Figure Supplement legends

**Figure 1-Figure Supplement 1. No evidence for a relationship between AB branch length and gene evolutionary rate (average MAD root-to-tip distance).** We did not detect a significant correlation between AB length and rate (average MAD-root to tip distance) when using genes with a non-zero (<0.00001) AB length. $p = 0.2025$, R =0.1143076, Pearson's correlation.

**Figure 1-Figure Supplement 2**. **No evidence for a relationship between AB branch length and gene evolutionary rate (average MAD root-to-tip distance).** The analysis is the same as in Figure 1-Figure Supplement 3, but with the inclusion of markers where AB branch length ~0. $p = 0.0761$, R = 0.102226, Pearson's correlation.

**Figure 1-Figure Supplement 3. A significant positive relationship between relative AB distance and evolutionary rate (MAD root-to-tip distance).** Genes with a greater relative AB distance have moderately higher evolutionary rates. $p = 0.007435$, R = 0.1537479, Pearson's correlation.

**Figure 1-Figure Supplement 4. Two proxies for marker gene verticality, ΔLL and between-domain split score, are highly correlated.** $p < 2.2 \times 10^{-16}$, R = 0.836679. Pearson's correlation.

**Figure 1-Figure Supplement 5. Low-verticality genes (as measured by ΔLL) have a higher evolutionary rate (as measured by mean root-to-tip distance on MAD-rooted gene trees).** Less vertically-evolving marker genes evolve faster, although the effect is moderate ($p$ = 0.01506, R = 0.1397803, Pearson's correlation).

**Figure 1-Figure Supplement 6. Low-verticality genes (measured by between-domain split score) have a higher evolutionary rate (MAD root-to-tip distance).** There is a positive relationship between between-domain split score (higher split score denotes lower verticality) and gene evolutionary rate (measured as mean MAD root-to-tip distance) $p = 0.002947$, R = 0.1705415, Pearson's correlation.

**Figure 1-Figure Supplement 7. High-verticality marker genes have longer AB branch lengths.** The depicted relationship is negative because high values of ΔLL correspond to lower between-domain verticality. This is the same analysis as in Figure 1C, but with marker genes with AB branch length ~0 included. The trendline is estimated using LOESS regression. We used a LOESS regression as the trendline here as the relationship varies across markers of differing verticality, $p = 0.009013$, R = -0.2317894, Pearson's correlation.

**Figure 1-Figure Supplement 8. AB branch length and relative AB distance are positively correlated.** This is the same analysis as in Figure 1B, but with marker genes with AB branch length ~0 included, $p < 2.2 \times 10\text{-}16$, R =0.706099, Pearson's correlation.

**Figure 1-Figure Supplement 9. AB branch length is negatively correlated with between-domain split score.** This is the same analysis as Figure 1E but with marker genes with AB branch length ~0 included, $p = 1.111 \times 10\text{-}05$, R = -0.2498829, Pearson's correlation.

**Figure 1-Figure Supplement 10. Within-domain split score and ΔLL are strongly correlated, suggesting that both proxies capture a common signal of marker gene verticality.** $p$ < 2.2×10-16, R = 0.6201967, Pearson's correlation.

**Figure 1-Figure Supplement 11. Low-verticality marker genes (measured as within-domain split score) have shorter relative AB distances.** A higher split score denotes lower verticality. $p$ = 5.685×10-06, R = -0.2577625, Pearson's correlation.

**Figure 1-Figure Supplement 12. Low-verticality marker genes (measured as within-domain split score) have shorter AB branch lengths**. A higher split score denotes lower verticality. $p$ = 0.0001467, R = -0.3318924, Pearson's correlation

**Figure 1-Figure Supplement 13. Low-verticality marker genes (measured as within-domain split score) have shorter AB branch lengths.** This is the same analysis as in Figure 1-Figure Supplement 12, but with marker genes with AB branch length ~0 included. There is a significant, but weaker negative correlation between AB branch length and marker gene verticality (within-domain split score) compared to the analysis where this class of genes is excluded. $p$ = 7.498×10-06, R = -0.2545369, Pearson's correlation.

**Figure 1-Figure Supplement 14**. **Raw count and percentage distribution of GTDB-defined classes for 10,575 archaeal and bacterial genomes in the expanded marker set analysis.**

**Figure 1-Figure Supplement 15**. **Raw count and percentage distribution of GTDB-defined phyla for 10,575 archaeal and bacterial genomes in the expanded marker set analysis.**

**Figure 1-Figure Supplement 16**. **Raw count and percentage distribution of domains for 10,575 archaeal and bacterial genomes in the expanded marker set analysis.**

**Figure 2-Figure Supplement 1. Among vertically-evolving marker genes, the split scores of ribosomal and non-ribosomal proteins are statistically indistinguishable.** The plot shows functional classification of markers (ribosomal markers or other) against the split score (a higher split score denotes greater disagreement with established within-domain relationships) using 54 markers from the new analysis. After removing genes that do not appear to have been vertically inherited since the divergence of Archaea and Bacteria, split scores of ribosomal and non-ribosomal markers were statistically indistinguishable (p = 0.8275, Wilcoxon rank sum test).

**Figure 3-Figure Supplement 1. Slow- and fast-evolving sites support different shapes for the universal tree.** (i) Tree of Archaea (blue) and Bacteria (red) inferred from a concatenation of 27 core genes using the best-fitting model (LG+C60+G4+F); (ii) Tree inferred from the fastest-evolving sites; (iii) Tree inferred from the slowest-evolving sites**.** Identical scale bars are provided for comparison.

**Figure 3-Figure Supplement 2. Vertically-evolving genes and slow-evolving sites support a longer relative AB branch length.** We estimated site-specific evolutionary rates for all marker genes in the expanded dataset (A-B), as well as for the 20 genes with the smallest ΔLL (top 5%) in that dataset (C-D). Concatenations based on the slowest sites (A,C) and on the top

5% vertical genes (C,D) support a longer AB branch. This suggests that the inference of a short AB branch is impacted by both substitutional saturation and unmodelled interdomain transfer of marker genes. Phylogenies were inferred under the LG+G4+F model in IQ-TREE 2. Branch lengths are the expected number of substitutions per site, as indicated by the scale bars. Alignment lengths in amino acids: A: 36797, B: 67274, C: 2736, D: 3884.

**Figure 3-Figure Supplement 3. The effect of modelling site compositional heterogeneity on AB branch length.** Increasing the number of protein mixture profiles, as well as trimming poorly-aligned positions, is associated with a change in AB branch length on the expanded marker set. All analyses used LG exchangeabilities, four rate categories (Gamma-distributed or freely estimated), and included a general composition vector containing the empirical amino acid frequencies (+F). Modelling of site heterogeneity with the C10-C60 models increases the inferred AB branch length ~2-fold. Trimming poorly-aligned sites slightly increases the AB branch estimation whereas relaxing the gamma rate categories slightly decreases estimation of AB branch length. LG (LG substitution matrix), G (four gamma rate categories), F (empirical site frequencies estimated from the data), C10-60 (number of protein mixture profiles used) R (four free rate categories which relax the assumption of a gamma distribution for rates, BMGE (trimming using Block Mapping and Gathering with Entropy)(Criscuolo and Gribaldo, 2010).

**Figure 4-Figure Supplement 1.** Raw count and percentage distribution of GTDB-defined classes of 350 archaea and 350 bacteria used in the 27 marker gene set analysis.

**Figure 4-Figure Supplement 2.** Raw count and percentage distribution of GTDB-defined phyla of 350 archaea and 350 bacteria used in the 27 marker gene set analysis.

### Supplementary Files

**Supplementary File 1.xlsx** – Marker metadata, KO and Pfam annotations and descriptions, and manual inspection notes for reciprocal monophyly and presence of paralogues, for 381 marker genes used in Zhu et al., 2019 (Methods) (Zhu et al., 2019).

**Supplementary File 2.xlsx** – Marker metadata, KO and Pfam annotations and descriptions, and manual inspection notes for 95 markers in the Core, Bacterial, and Non-Ribosomal marker gene sets (Methods).

**Supplementary File 3.xlsx** – NCBI taxonomic information for 350 Archaeal and 350 Bacterial genomes sampled in the new analyses

**Supplementary File 4.xlsx** – Clade definitions for quantifying taxonomic splits and split-score statistical summaries for ranking of the Core, Bacterial, Non-Ribosomal marker genes, and 381 marker genes (Methods).

**Supplementary File 5a** – Functional annotations for the top 20 genes used and in Figure 6 and referred to in Figure 7Aii of the main text.

**Supplementary File 5b. A list of fossil calibrations employed in relaxed molecular clock analyses**. All calibrations were modelled as uniform distributions between a hard minimum and a soft maximum. The probability that the maximum could be exceeded was modelled as a 2.5% probability tail.
