## Supplementary material for "An estimate of the deepest branches of the tree of life from ancient vertically-evolving genes": Figure 4-figure supplement 1

Class Distribution (Counts) of 700 ArcBac Reference Genomes

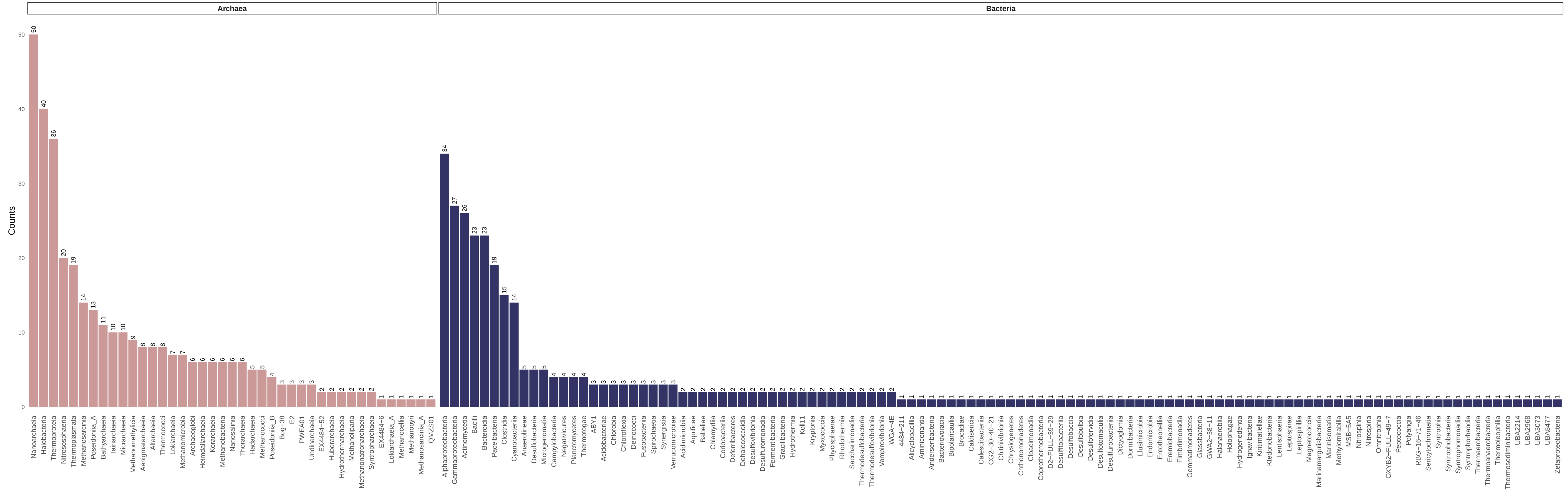


Class Distribution (Percentage) of 700 ArcBac Reference Genomes

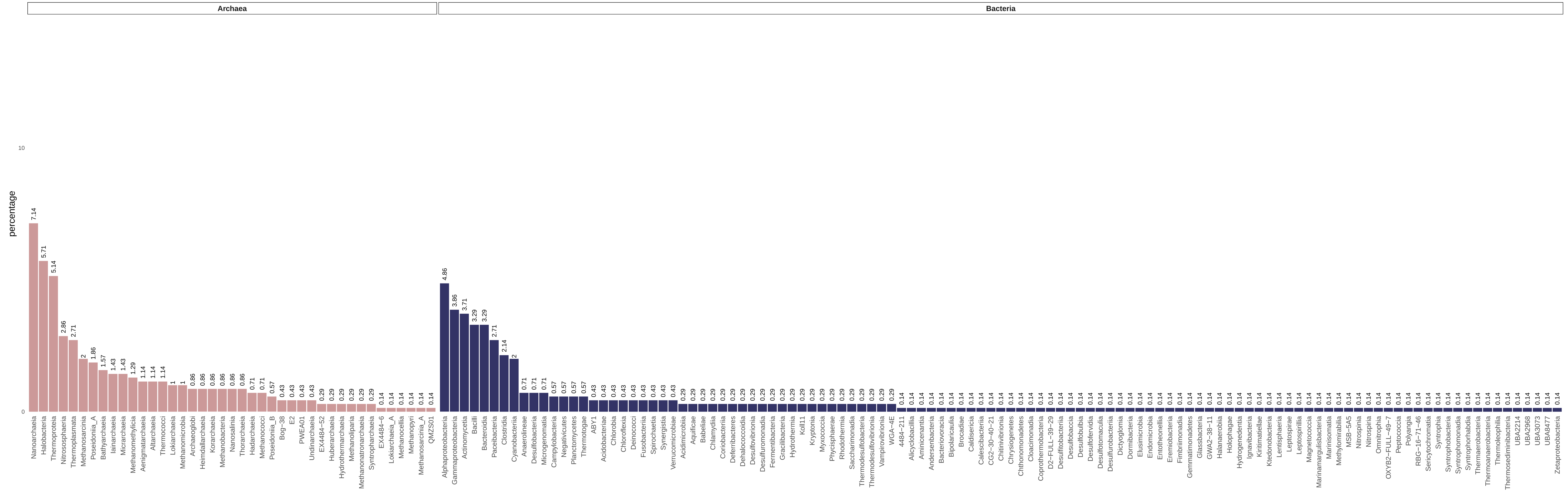
