## Supplementary figures and images for "An estimate of the deepest branches of the tree of life from ancient vertically-evolving genes"

### Figure 1-figure supplement 1

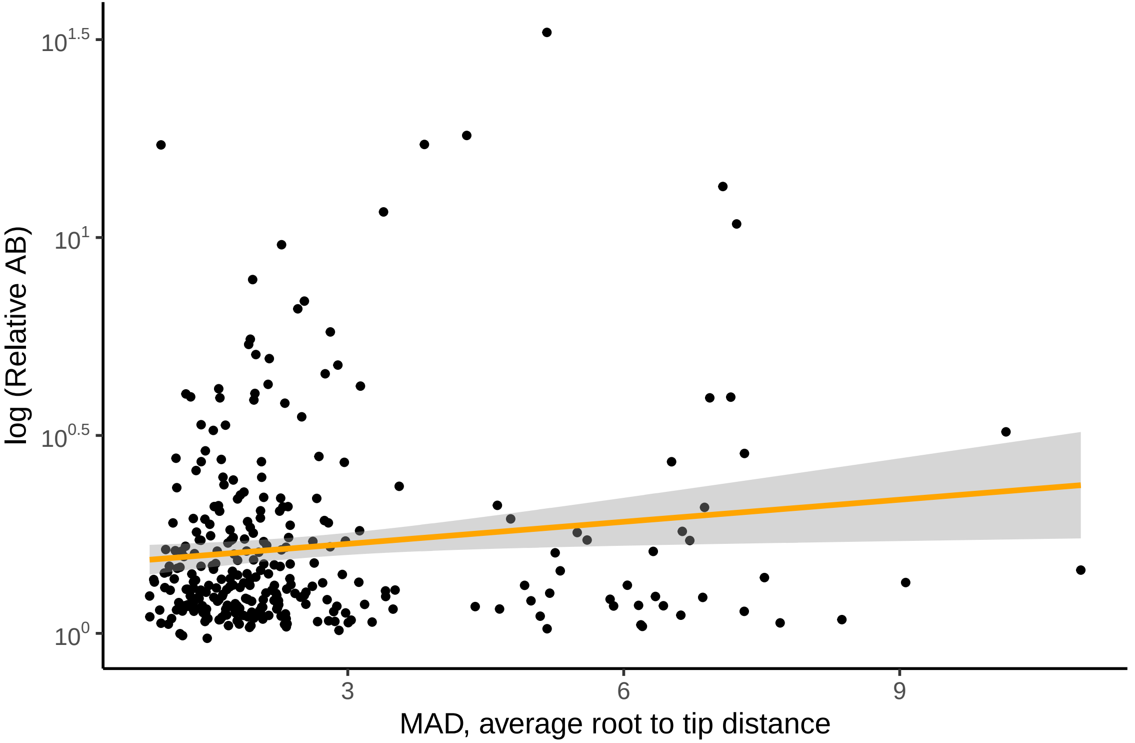

### Figure 1-figure supplement 2

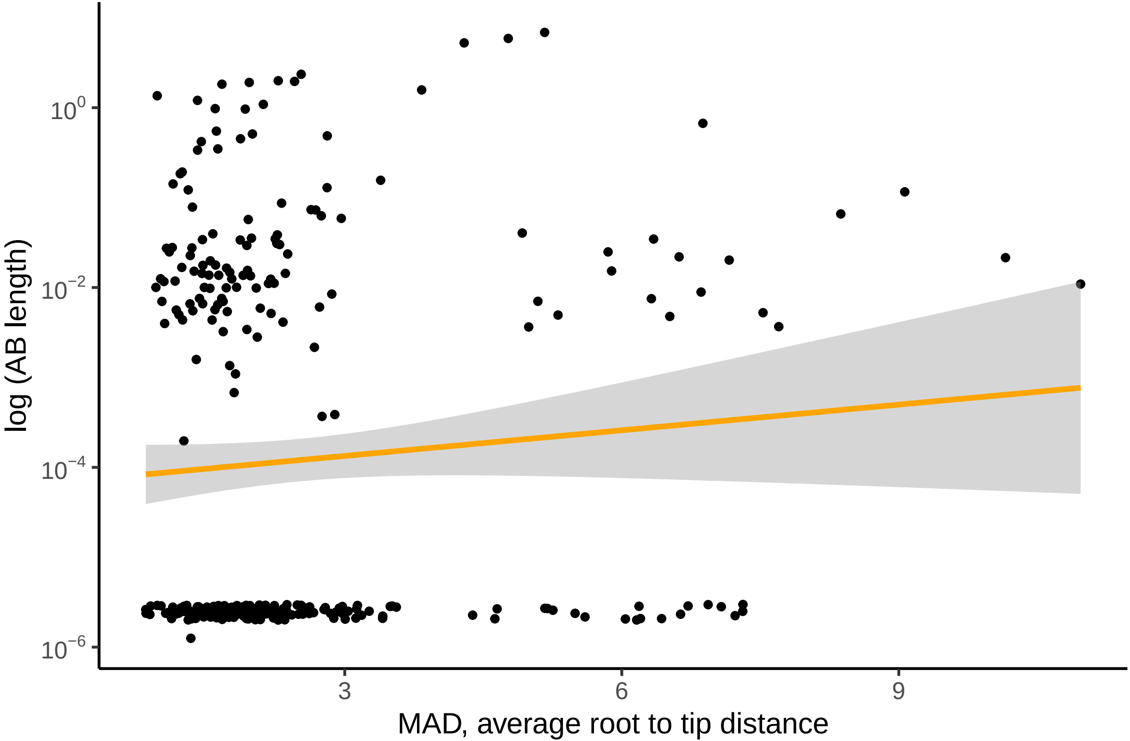

### Figure 1-figure supplement 3

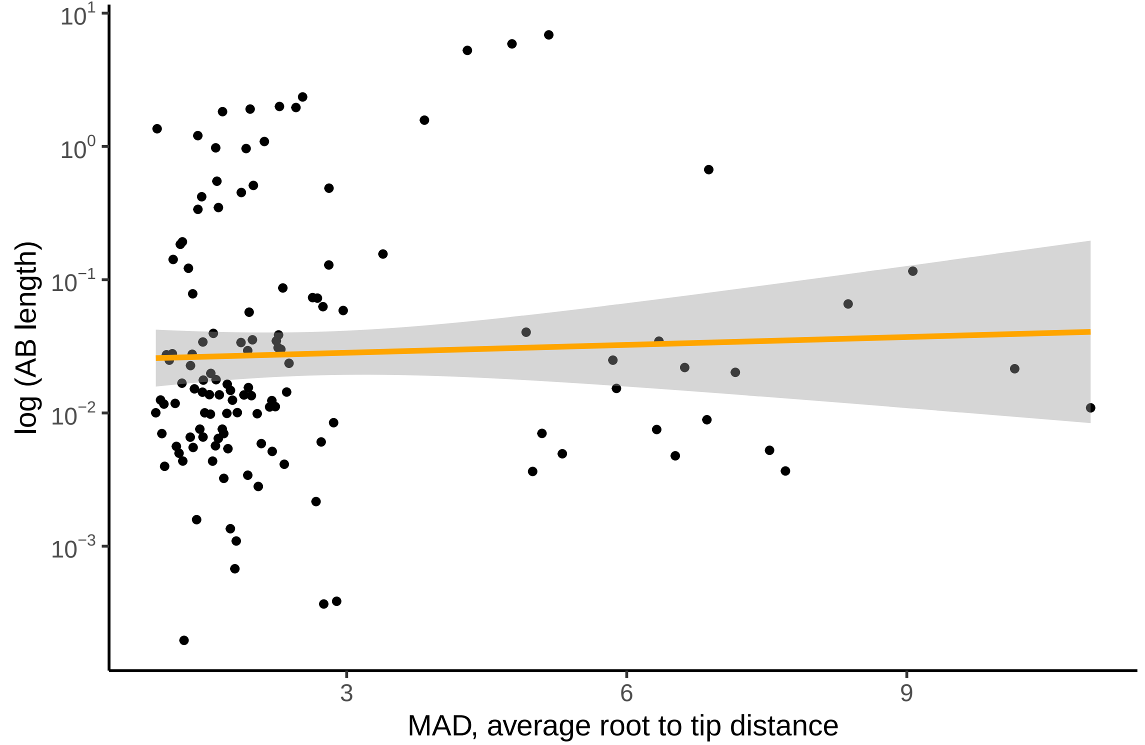

### Figure 1-figure supplement 4

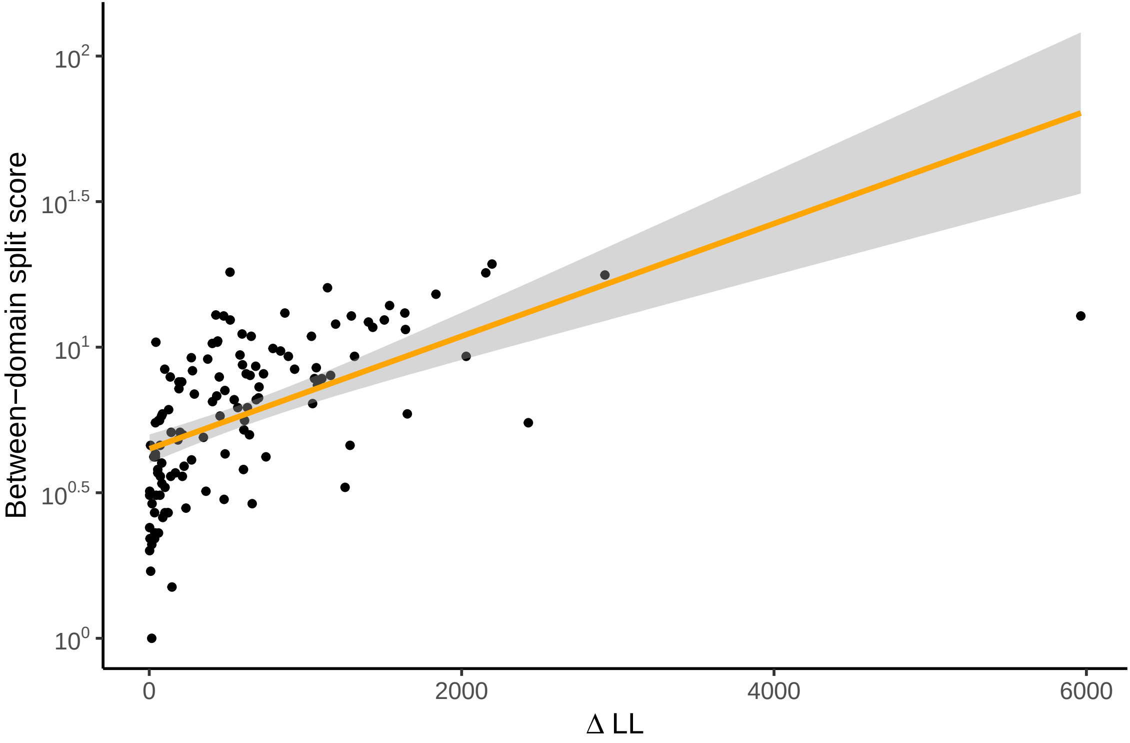

### Figure 1-figure supplement 5

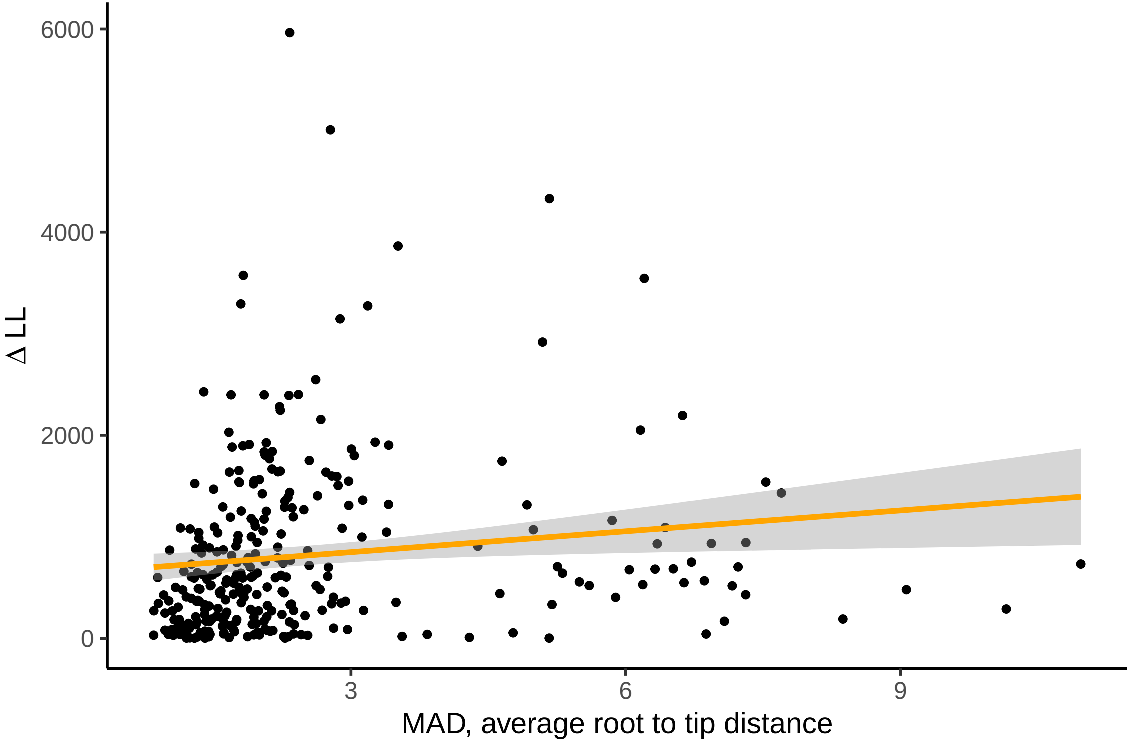

### Figure 1-figure supplement 6

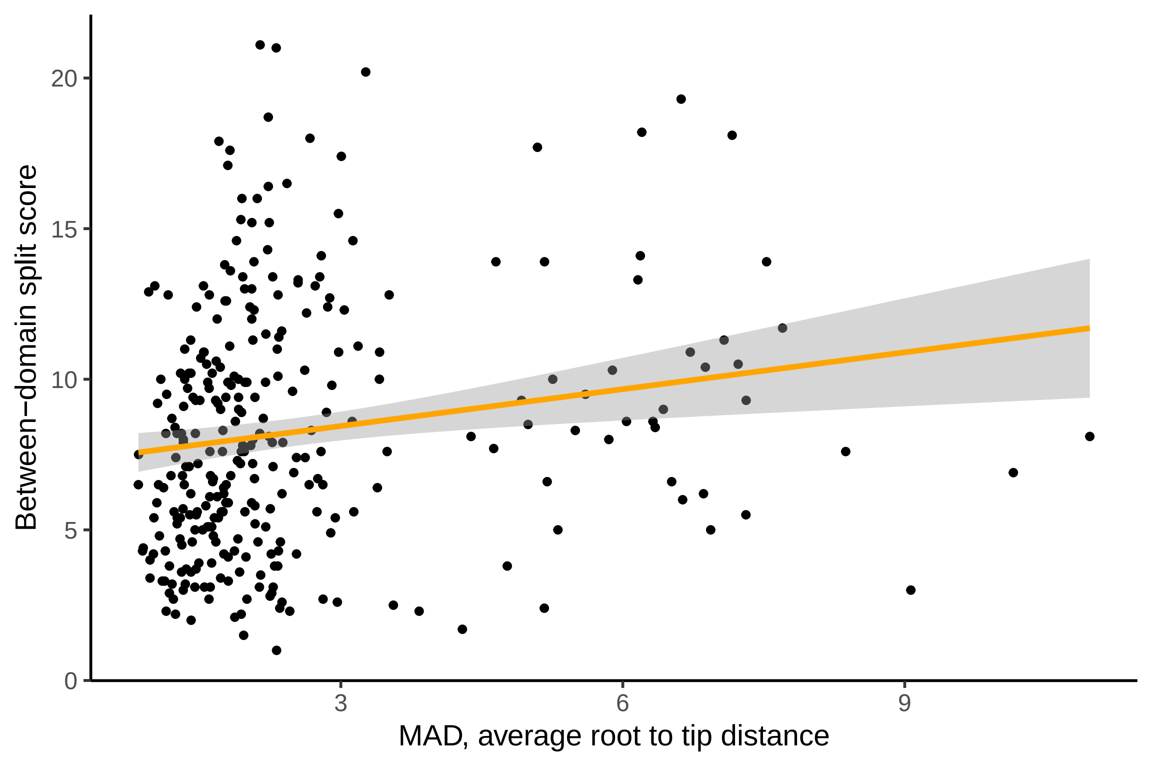

### Figure 1-figure supplement 7

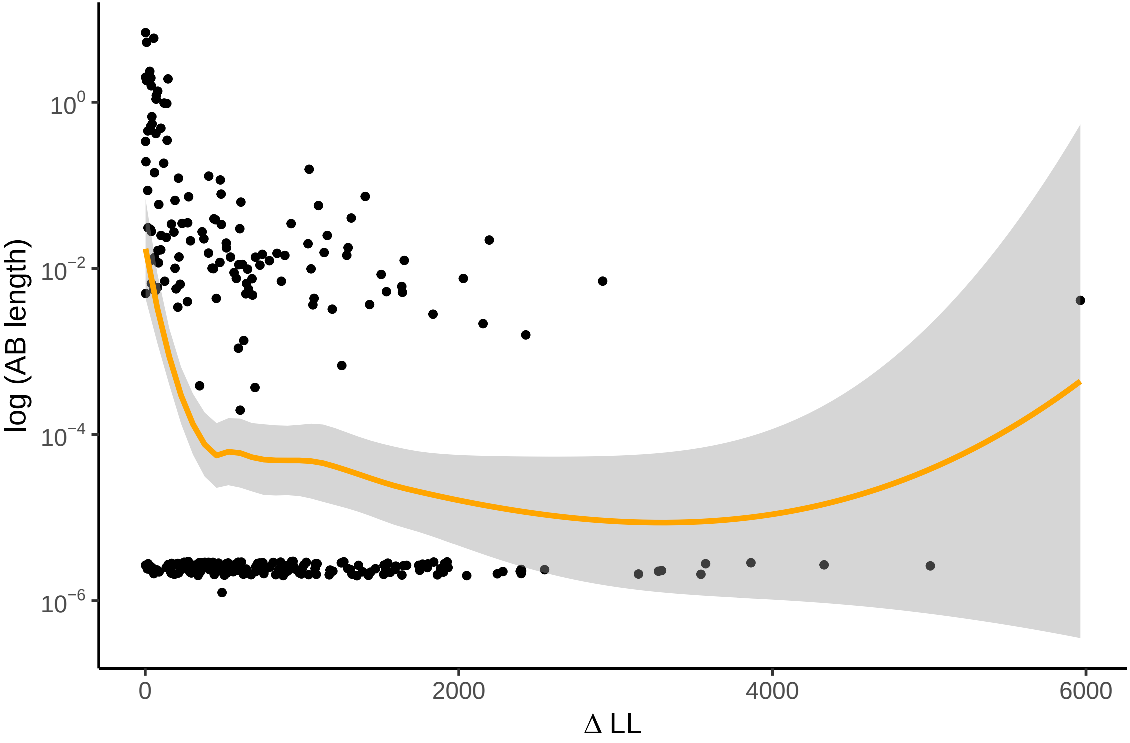

### Figure 1-figure supplement 8

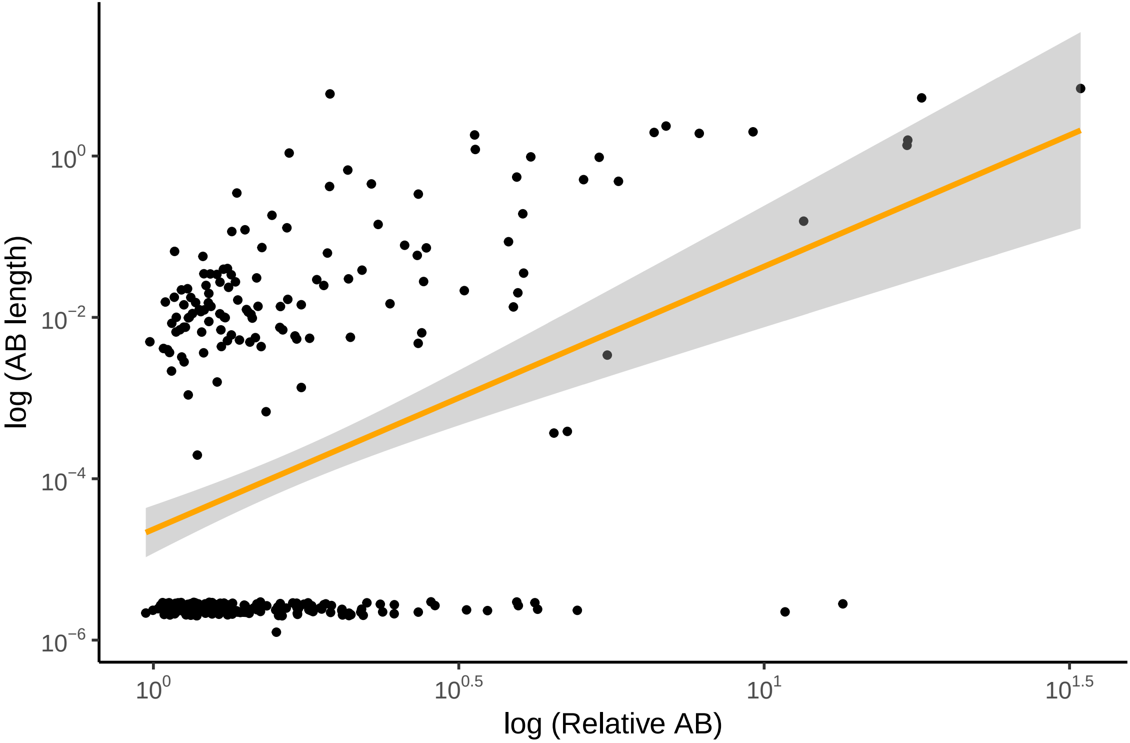

### Figure 1-figure supplement 9

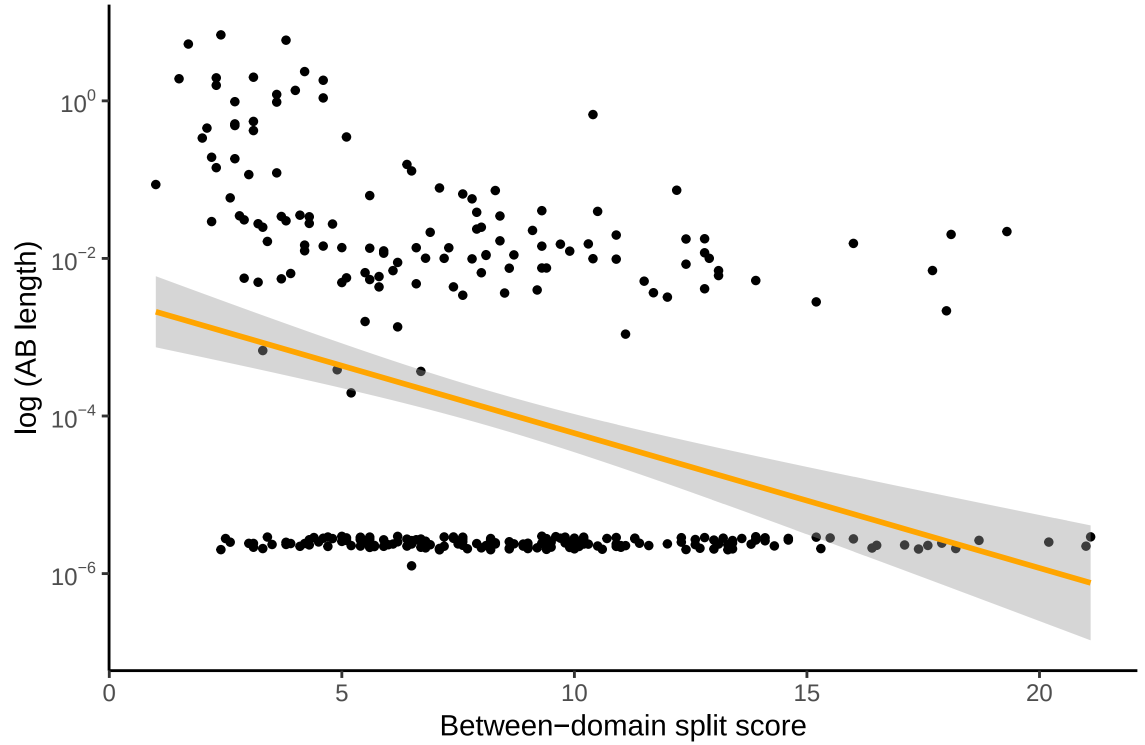

### Figure 1-figure supplement 10

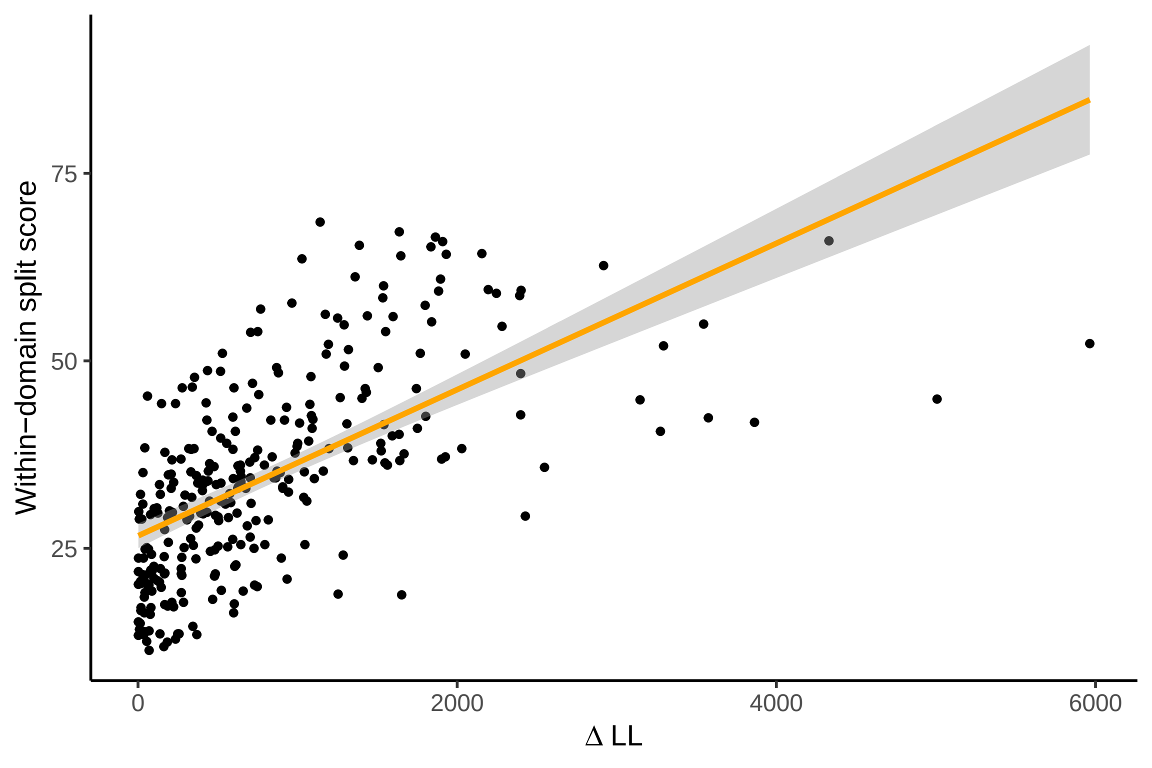

### Figure 1-figure supplement 11

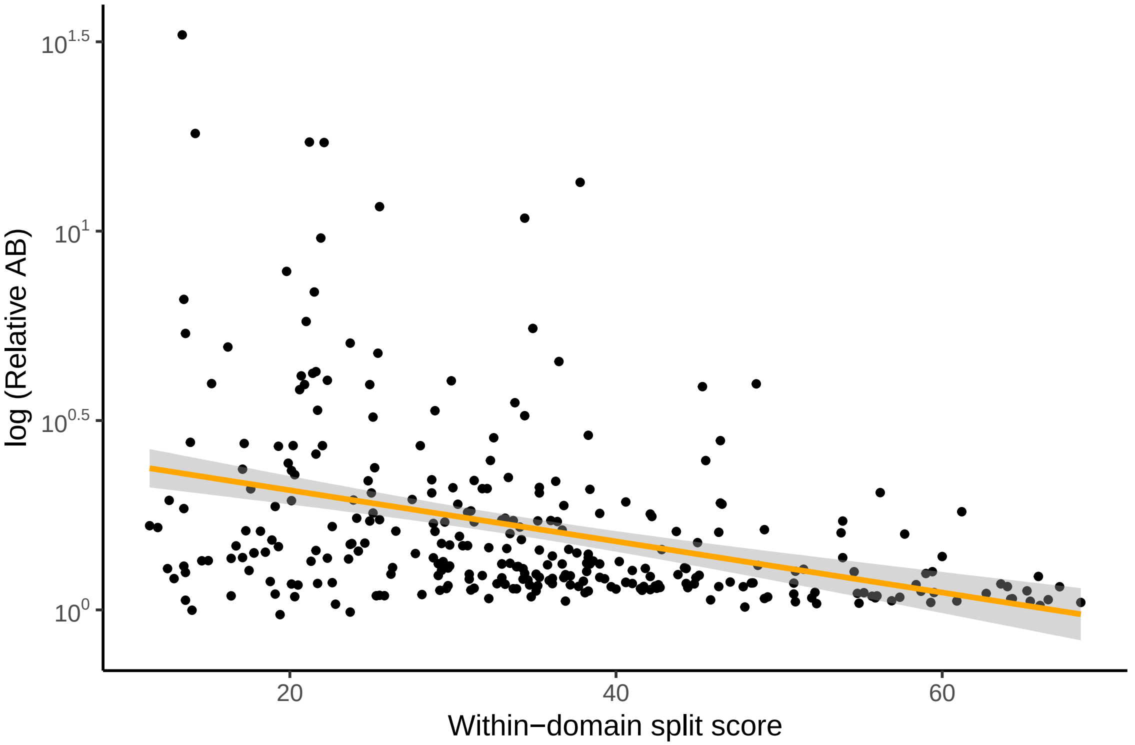

### Figure 1-figure supplement 12

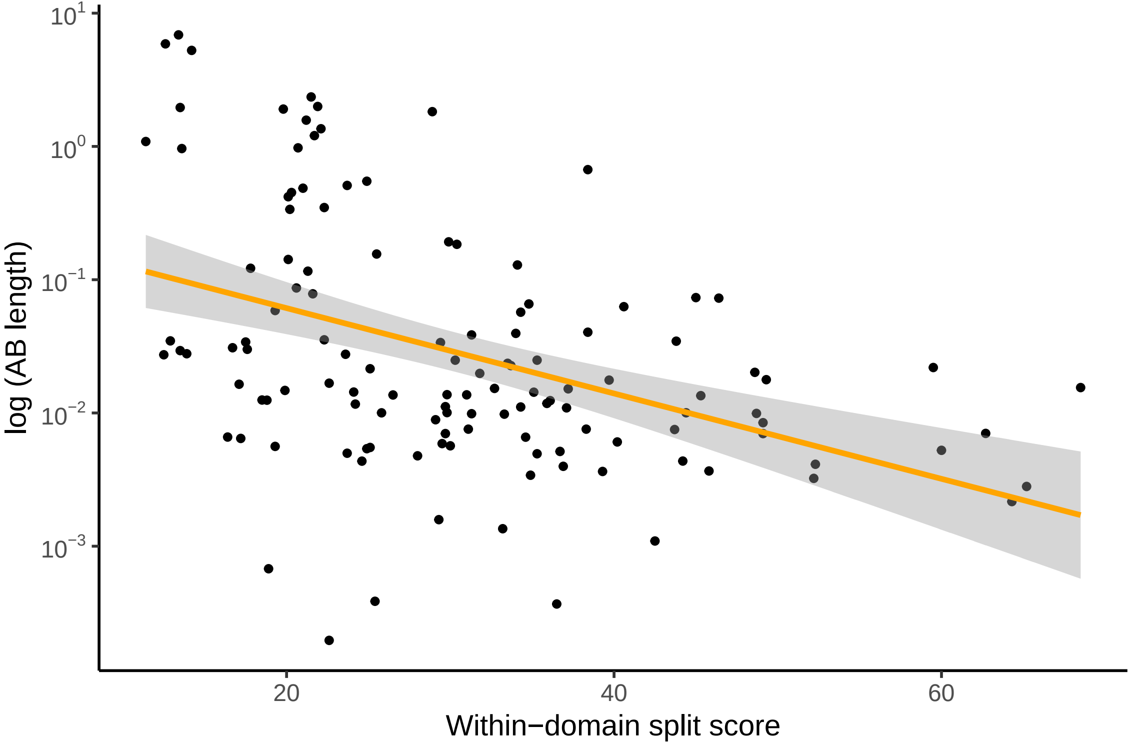

### Figure 1-figure supplement 13

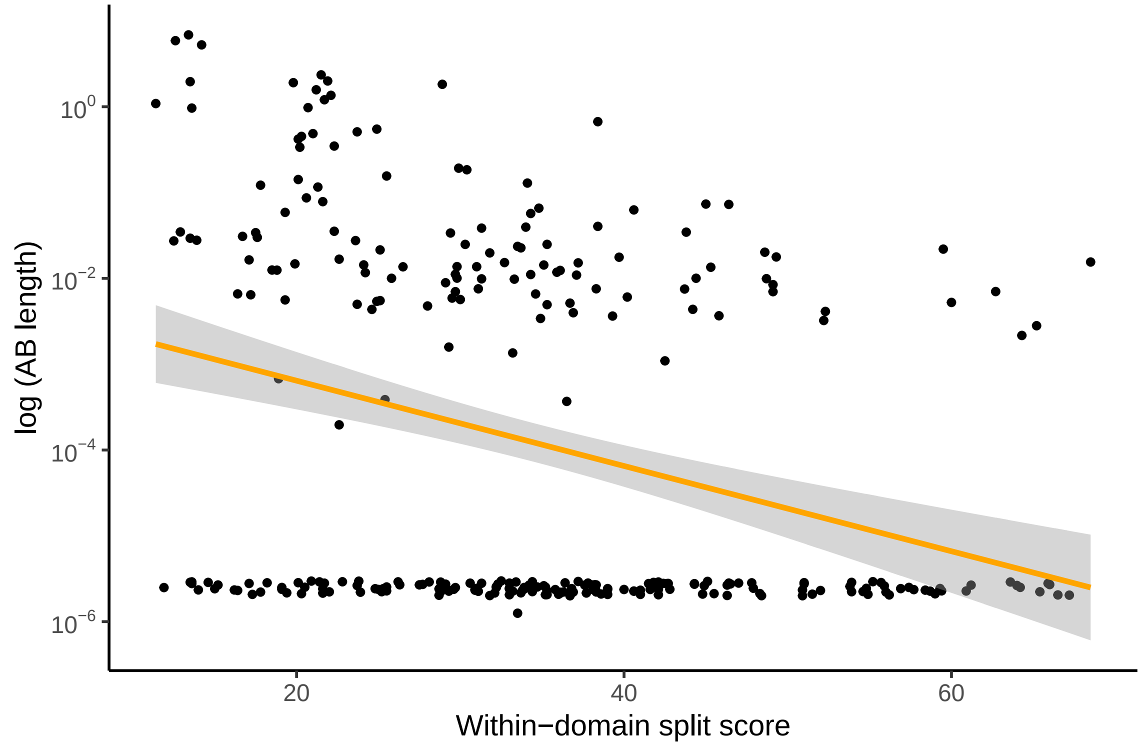

### Figure 1-figure supplement 14

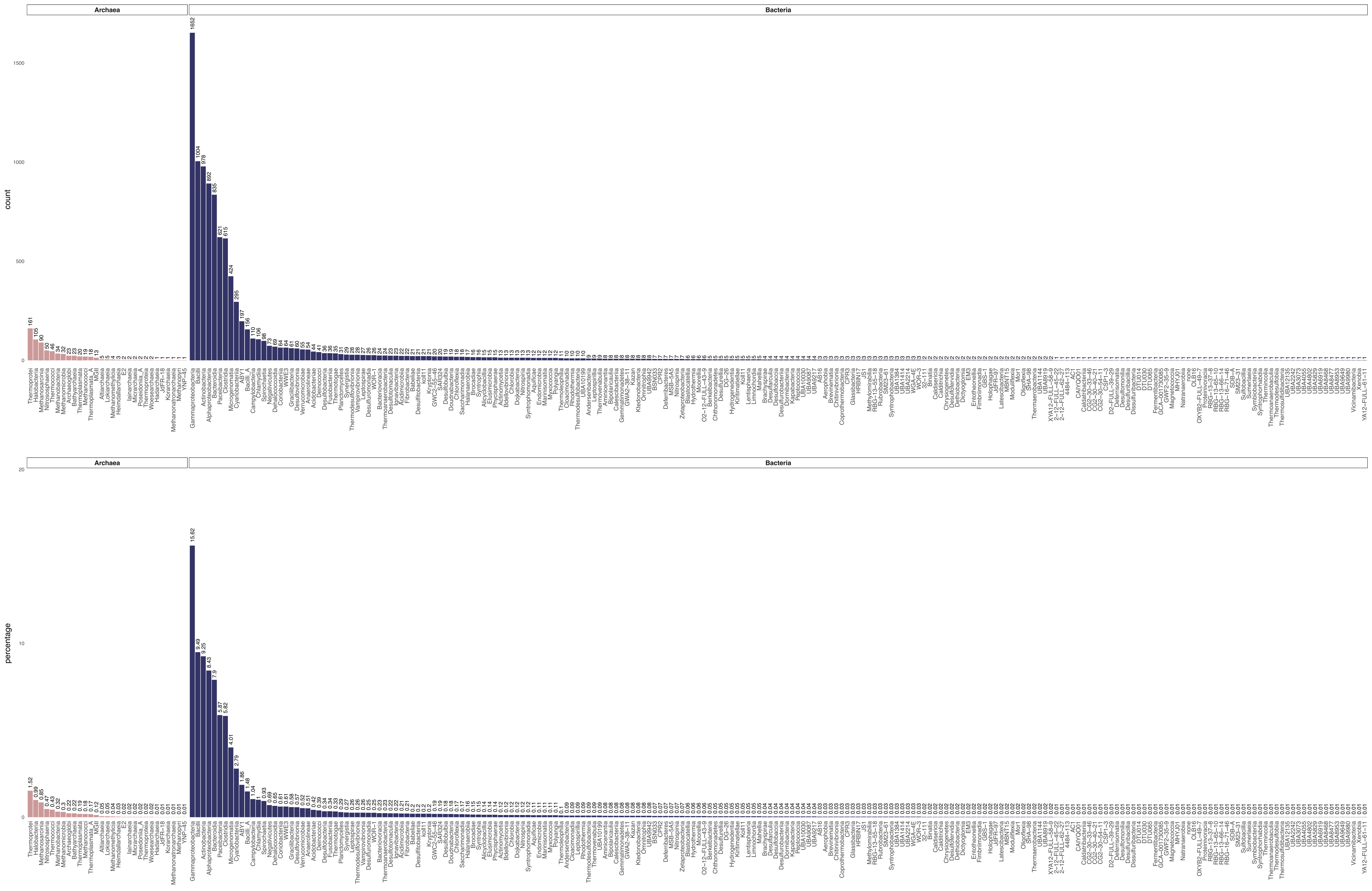

### Figure 1-figure supplement 15

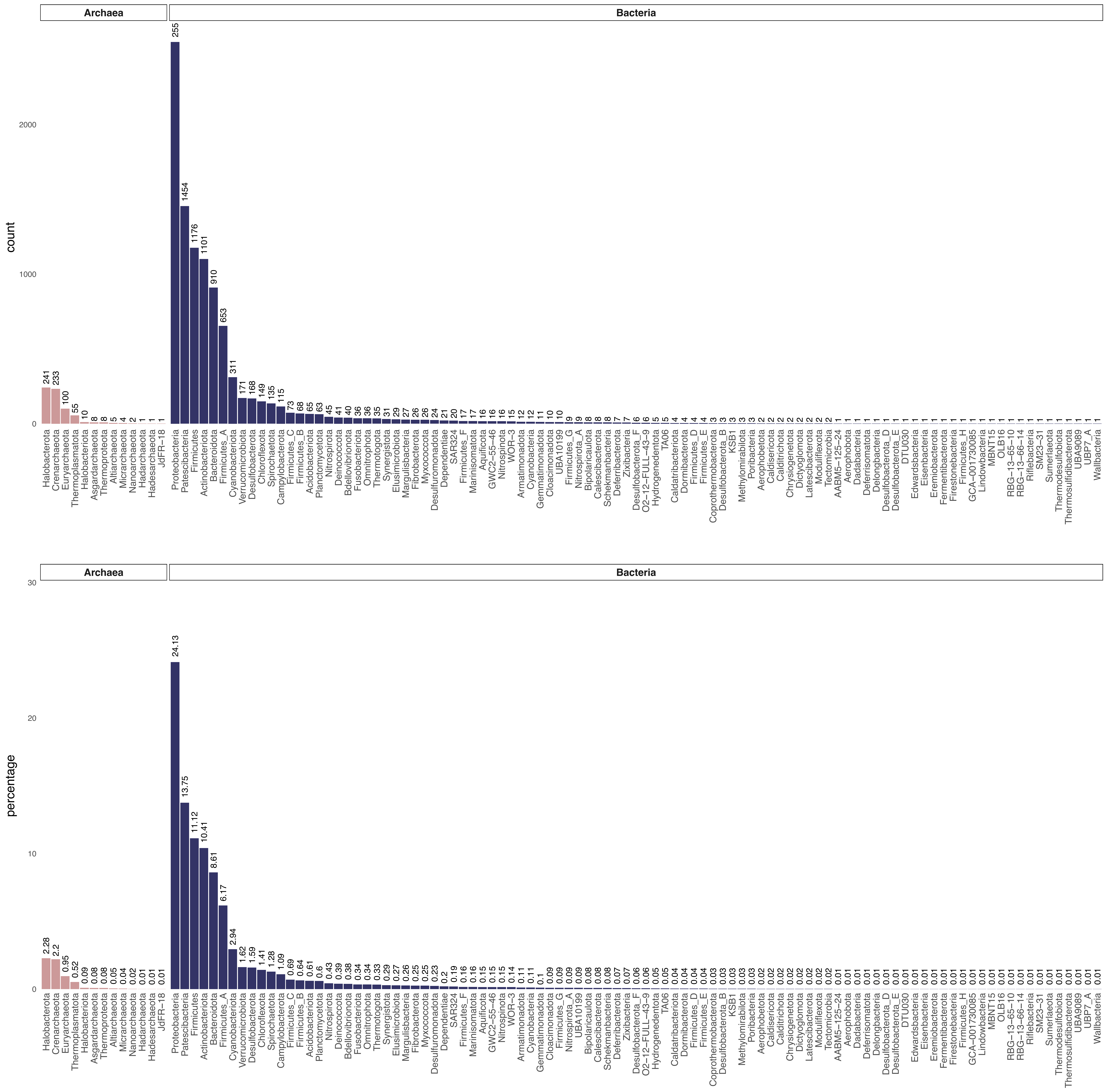

### Figure 1-figure supplement 16

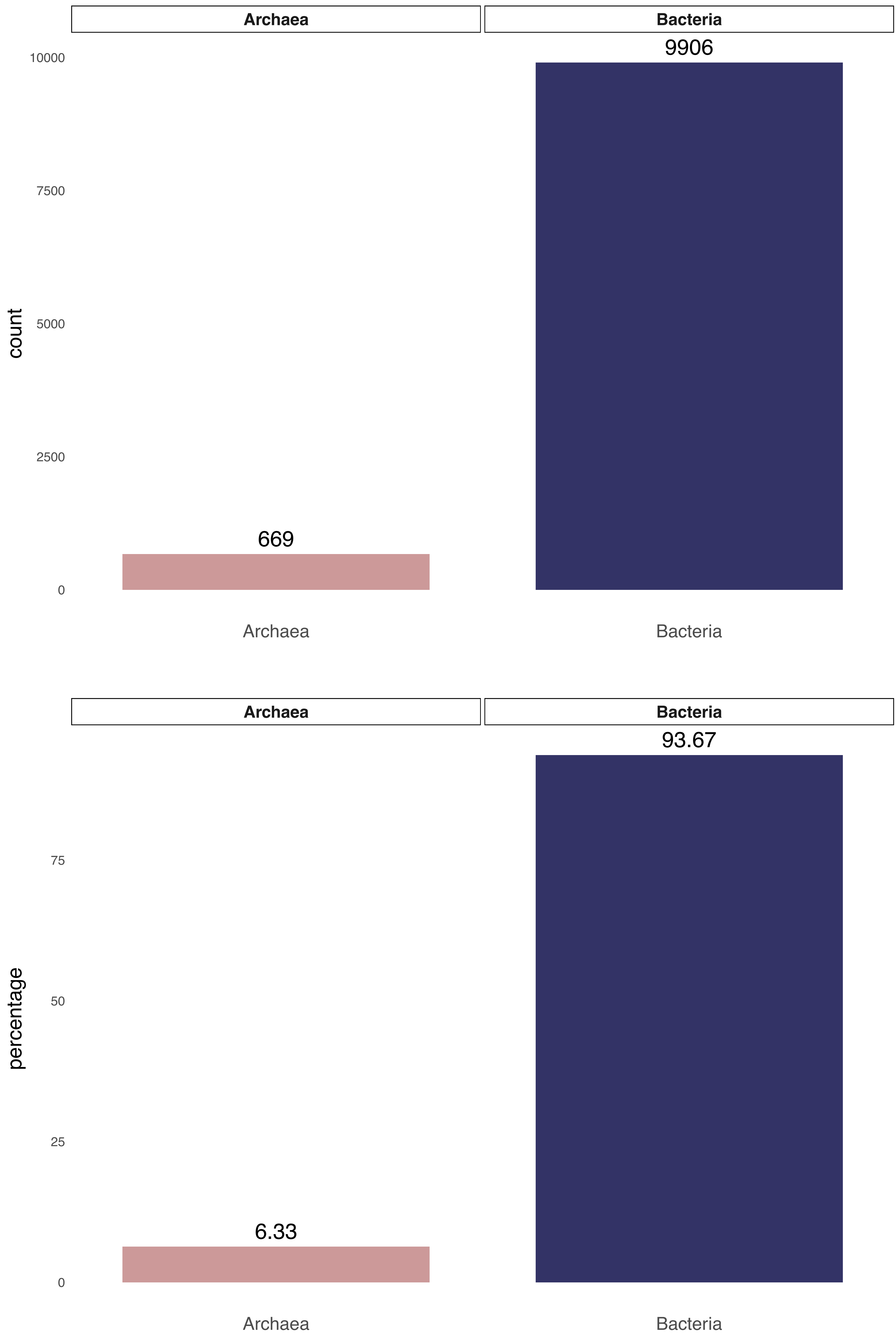

### Figure 2-figure supplement 1

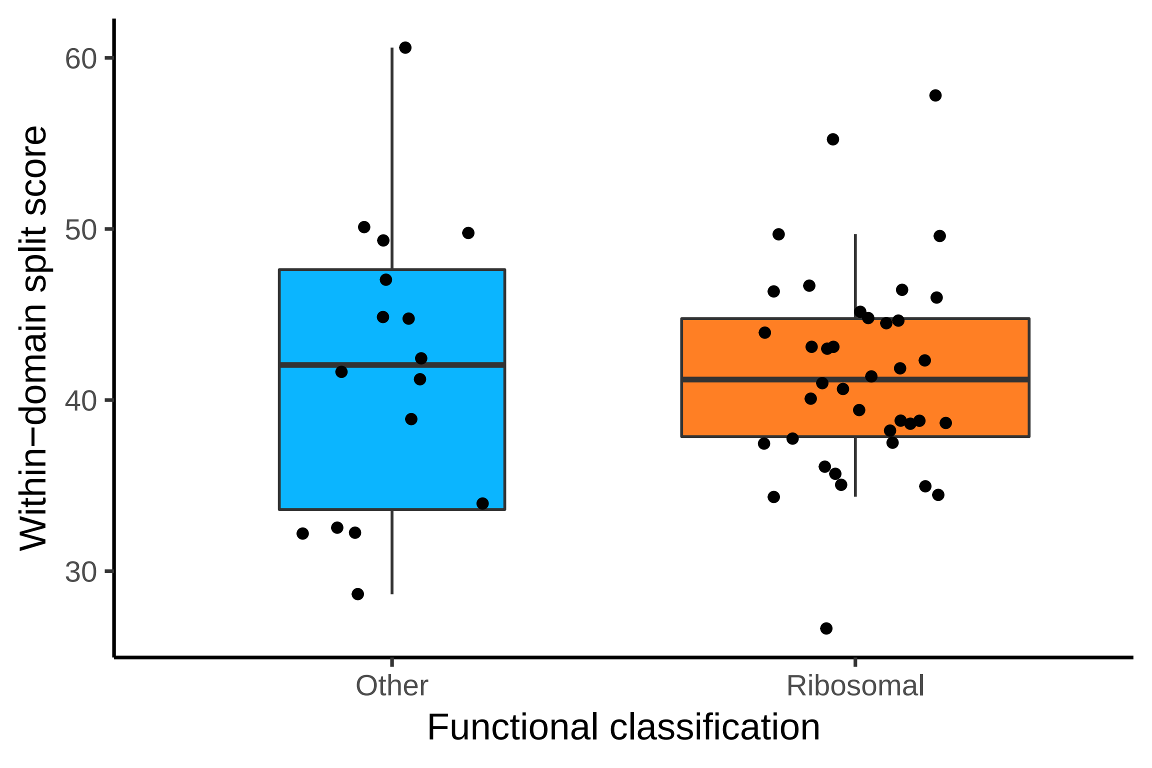

### Figure 3-figure supplement 1

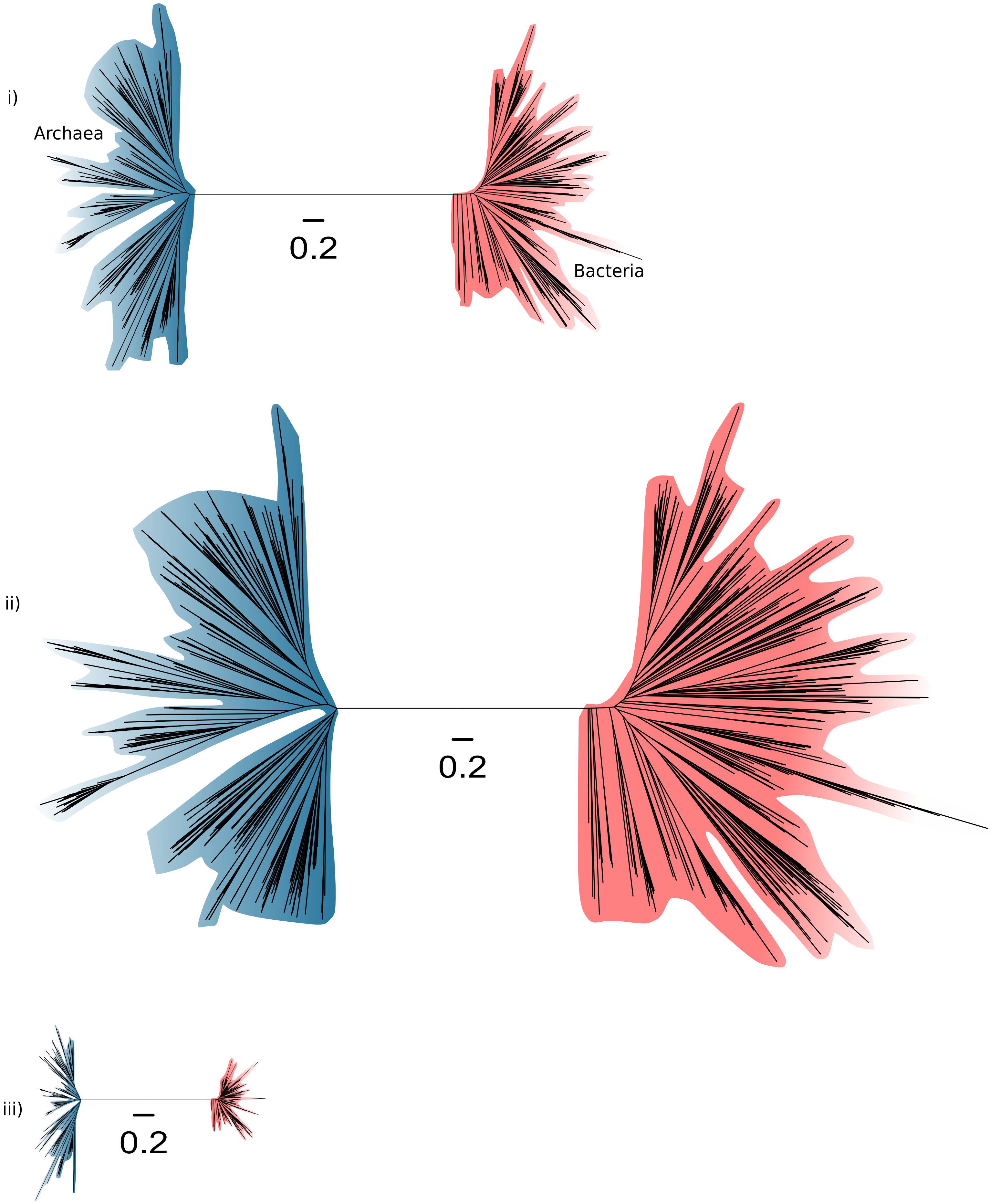

### Figure 3-figure supplement 2

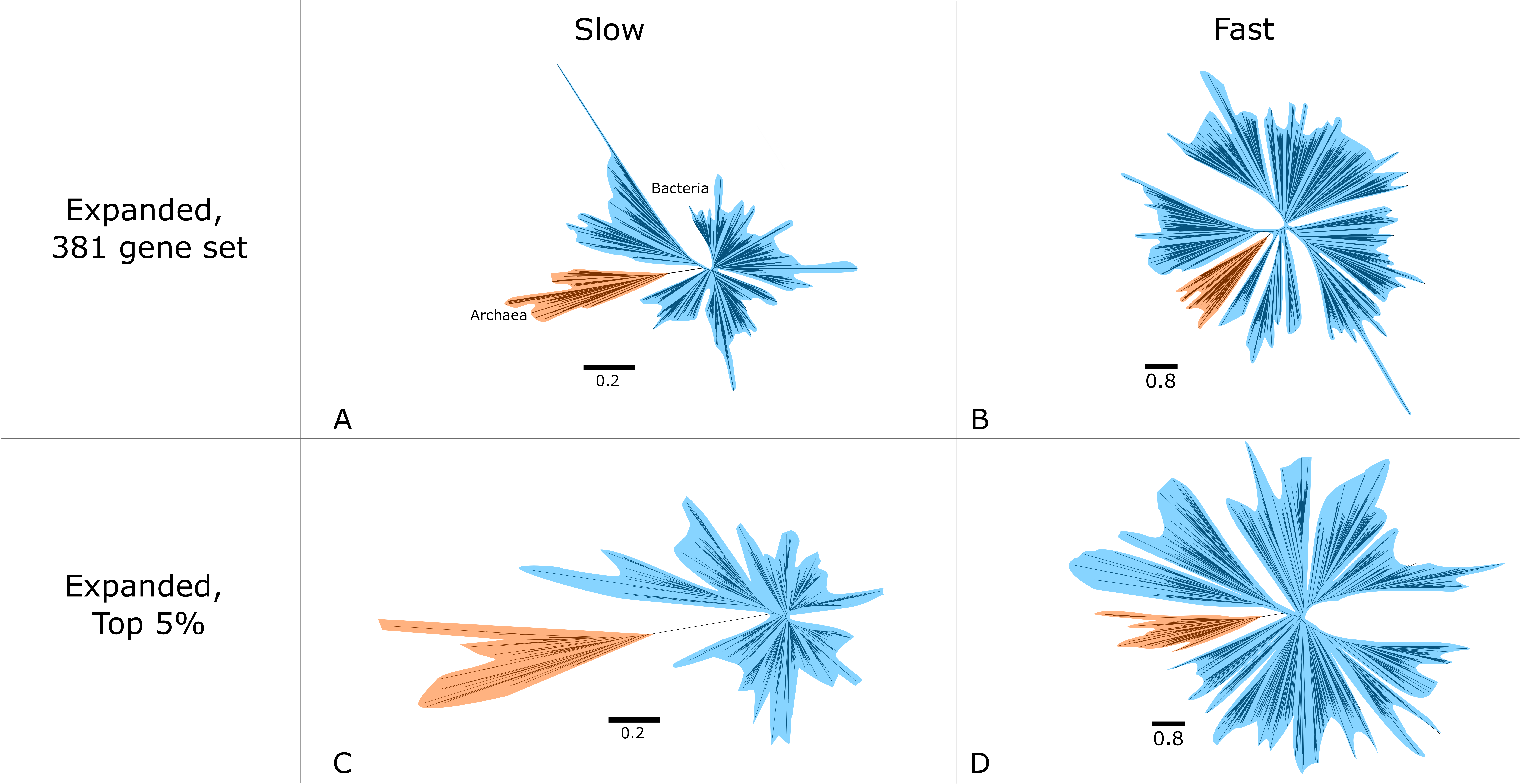

### Figure 3-figure supplement 3

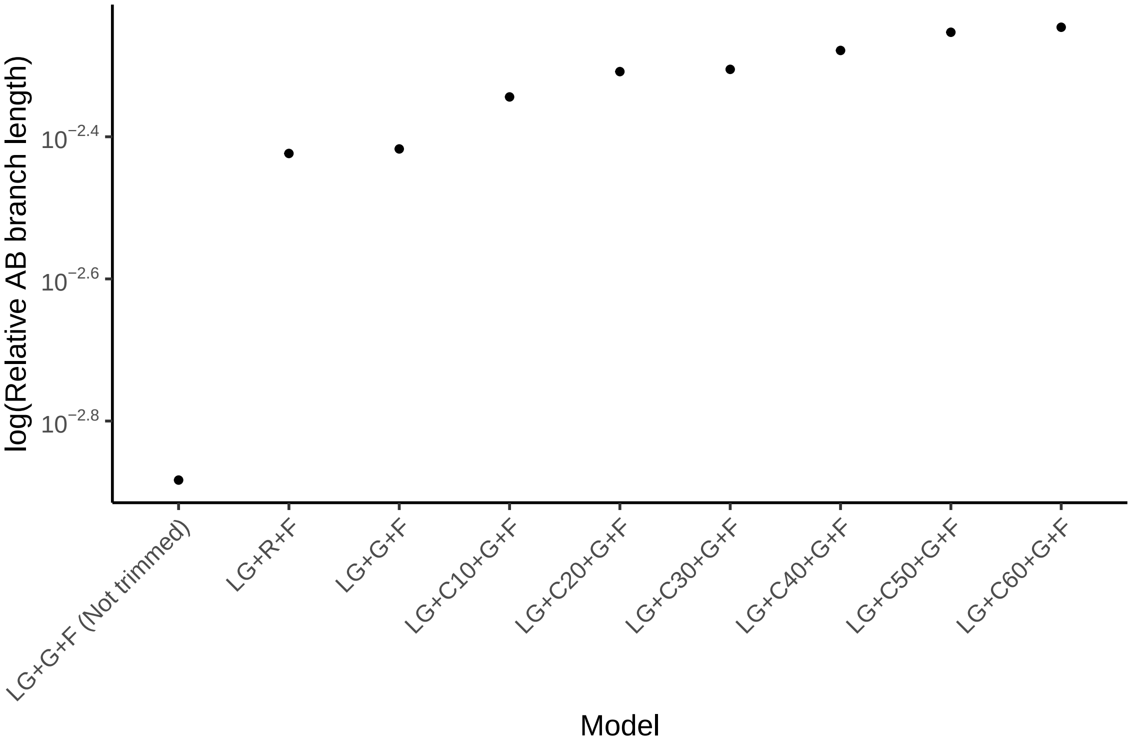
