## Supplementary File 5 for "An estimate of the deepest branches of the tree of life from ancient vertically-evolving genes"

**5a**

| **Expanded Marker** | **uniprot_id** | **Protein** | **Gene** |
| --- | --- | --- | --- |
| p0023 | D3R080 | Translation initiation factor IF-2 | infB |
| p0313 | Q2JK96 | Arginine--tRNA ligase (EC 6.1.1.19) (Arginyl-tRNA synthetase) (ArgRS) | argS |
| p0076 | D5SWY7 | tRNA-2-methylthio-N(6)-dimethylallyladenosine synthase (EC 2.8.4.3) ((Dimethylallyl)adenosine tRNA methylthiotransferase MiaB) (tRNA-i(6)A37 methylthiotransferase) | miaB |
| p0004 | P19486 | Elongation factor 1-alpha (EF-1-alpha) (Elongation factor Tu) (EF-Tu) | tuf |
| p0383 | E7H5S9 | Phosphoserine aminotransferase (EC 2.6.1.52) (Phosphohydroxythreonine aminotransferase) (PSAT) | serC |
| p0389 | P21469 | 30S ribosomal protein S7 (BS7) | rpsG |
| p0072 | Q2GDY4 | Phenylalanine--tRNA ligase beta subunit (EC 6.1.1.20) (Phenylalanyl-tRNA synthetase beta subunit) (PheRS) | pheT |
| p0010 | C9RQN5 | DNA-directed RNA polymerase subunit beta (RNAP subunit beta) (EC 2.7.7.6) (RNA polymerase subunit beta) (Transcriptase subunit beta) | rpoB |
| p0229 | E6SH25 | Arginine biosynthesis bifunctional protein ArgJ [Cleaved into: Arginine biosynthesis bifunctional protein ArgJ alpha chain; Arginine biosynthesis bifunctional protein ArgJ beta chain] [Includes: Amino-acid acetyltransferase (EC 2.3.1.1) (N-acetylglutamate synthase) (AGSase); Glutamate N-acetyltransferase (EC 2.3.1.35) (Ornithine transacetylase) (Ornithine acetyltransferase) (OATase)] | argJ |
| p0268 | E6SJH8 | Aspartate carbamoyltransferase (EC 2.1.3.2) (Aspartate transcarbamylase) (ATCase) | pyrB |
| p0027 | A9WK84 | Leucine--tRNA ligase (EC 6.1.1.4) (Leucyl-tRNA synthetase) (LeuRS) | leuS |
| p0182 | A0A0E8TCT0 | Glutamate synthase (EC 1.4.1.14) (EC 1.4.7.1) | gltB_1 |
| p0109 | Q4FV41 | DNA mismatch repair protein MutS | mutS |
| p0346 | B0VFP4 | Thymidine phosphorylase (EC 2.4.2.4) | deoA |
| p0162 | D3R256 | Ribosome biogenesis GTPase Era | era |
| p0030 | D9PW62 | ATP-dependent DNA helicase (EC 3.6.1.-) | pcrA1 |
| p0138 | C6D886 | Lon protease (EC 3.4.21.53) (ATP-dependent protease La) | lon |
| p0241 | E8R3Q8 | Glutamate 5-kinase (EC 2.7.2.11) (Gamma-glutamyl kinase) (GK) | proB |
| p0071 | N2BS08 | DNA polymerase I (EC 2.7.7.7) | polA |
| p0046 | B5YFL1 | Holliday junction ATP-dependent DNA helicase RuvB (EC 3.6.4.12) | ruvB |

**5b**

| Node | Fossil | Minimum | Soft Maximum | Reference |
| --- | --- | --- | --- | --- |
| 1. LUCA | Strelley Pool Formation, Western Australia | 3347 Ma | 4520 Ma | Betts et al. 2018 |
| 2. Total group Cyanobacteria | Manzimnyama Banded Ironstone Formation,  South Africa | 3225 Ma | 4520 Ma | Betts et al. 2018 |
| 3. Crown group Cyanobacteria | *Bangiomorpha pubescens* | 1033 Ma | 4520 Ma | Betts et al. 2018 |
| 4. Heterocystous/akinete-forming Cyanobacteria (Cyanobacteria sections IV+V) | *Anhuthruix magna* | 720 Ma | 4520 Ma | This paper |
| 5. Alphaproteobacteria | *Bangiomorpha pubescens* | 1033 Ma | 4520 Ma | Betts et al. 2018 |
| 6. Anoxychlamydiales | *Bangiomorpha pubescens* | 1033 Ma | 4520 Ma | This paper |
| 7. Lokiarchaeota | Changzhougou Formation, Northern China | 1619 Ma | 4520 Ma | Betts et al. 2018 |

**Calibration 4**

***Fossil taxon and specimen:*** *Anhuthruix magna* [Llb x4B, Institute of Geology and Paleontology, Technical University of Berlin,Berlin, Germany] from the Liulaobei Formation in the Huainan region of North China

***Phylogenetic justification:*** *A. magna* was originally described as a tubular organism (Steiner 1994), however Pang et al. (2018) interpreted it as a filamentous cyanobacteria based on new material.  Based on the presence of binary fission, hormogonia, akinetes, and probably heterocysts, an affinity with subsections IV+V of cyanobacteria was proposed.

***Minimum age:*** 720 Ma

***Maximum age:*** 4520 Ma

***Age justification:*** The Liulaobei Formation forms part of the Huainan Group in Anhui, Northern China. Geochronometric data from the Huainan Group proposed a Tonian age (840 +/- 72 Ma; Wang et al. 1984), but these measures have since been determined as unreliable (Dong et al. 2008). A pre-Cryogenian age was proposed based on stromatolite biostratigraphy and the presence of *Chuaria*, *Ellipsophysa*, and *Tawuia* in the Huainan Group (Cao et al. 1985; Fu 1989). This is further supported by the presence of the acanthomorphic acritarch *Trachyhystrichosphaera aimika* in the Liulaobei Formation and an overall acritarch assemblage that is distinct from those of the Ediacaran (Tang et al. 2013). Thus, a minimum age based on the upper boundary of the Tonian, 720 Ma (Van Kranendonk et al. 2012).

**Calibration 6**

**Crown Chlamydiia/Chlamydiae/Anoxychlamydiales**

***Fossil taxon and specimen***: *Bangiomorpha pubescens*. (Holotype) HUPC 62912, Slide HUST-1A, England Finder coordinates: O-35.

***Locality and Stratigraphy level***: Lower Hunting Formation, Somerset Island, Arctic Canada. ***Soft Minimum age***: 1033 Ma (1092 Ma ± 59 Myr )

***Soft Maximum age***: 4520 Ma (4510 Ma ± 10 Myr)

***Description***: Recent work (Stairs et al. 2020) suggests that members of the Anoxychlamydiales (including a lineage named Chlam. Bact SM23_39), a clade within Chlamydiia (and Chlamydiae), donated at least three subunits of hydrogenase maturase (E, F, G) to stem eukaryotes prior to the radiation of crown eukaryotes. These phylogenies imply that a minimum age for Anoxychlamdiales is the age of the oldest crown eukaryote and thus a minimum age is derived following Betts et al. 2020

***References***

Betts, H.C., Puttick, M.N., Clark, J.W. *et al.* (2018) Integrated genomic and fossil evidence illuminates life’s early evolution and eukaryote origin. *Nat Ecol Evol* **2**1556–1562

Steiner, M. (1994) *Die neoproterozoischen Megaalgen südchinas*. Selbstverlag Fachbereich Geowissenschaften, 1994.lip

Pang, K., Tang, Q., Chen, L., Wan, B., Niu, C., Yuan, X., & Xiao, S. (2018). Nitrogen-fixing heterocystous cyanobacteria in the Tonian period. *Current Biology*, *28*(4), 616-622.

Dong, L., Xiao, S., Shen, B., Yuan, X., Yan, X., & Peng, Y. (2008). Restudy of the worm-like carbonaceous compression fossils Protoarenicola, Pararenicola, and Sinosabellidites from early Neoproterozoic successions in North China. *Palaeogeography, Palaeoclimatology, Palaeoecology*, *258*(3), 138-161.

Van Kranendonk, M. J., Altermann, W., Beard, B. L., Hoffman, P. F., Johnson, C. M., Kasting, J. F., Melezhik, V. A., Nutman, A. P., Papineau, D., & Pirajno, F. (2012). Chronostratigraphic division of the precambrian: Possibilities and challenges. In *The Geologic Time Scale 2012* (pp. 299-392). Elsevier B.V.

Cao, R., W. Zhao, and G. Xiao. "Late Precambrian stromatolites from north Anhui Province." *Mem. Nanjing Inst. Geol. Paleont. Acad. Sin* 21 (1985): 1-84.

Jun-hui, F. (1989). Assemblage of Huainan biota and its characteristics. *Acta Palaeontologica Sinica*, *5*.

Tang, Q., Pang, K., Xiao, S., Yuan, X., Ou, Z., & Wan, B. (2013). Organic-walled microfossils from the early Neoproterozoic Liulaobei Formation in the Huainan region of North China and their biostratigraphic significance. *Precambrian Research*, *236*, 157-181.

Stairs, C.W., Dharamshi, J.E., Tamarit, D., Eme, L., Jørgensen, S.L., Spang, A. and Ettema, T.J., 2020. Chlamydial contribution to anaerobic metabolism during eukaryotic evolution. *Science advances*, *6*(35), p.eabb7258.
